# Rhapsody: Pathogenicity prediction of human missense variants based on protein sequence, structure and dynamics

**DOI:** 10.1101/737429

**Authors:** Luca Ponzoni, Zoltán N. Oltvai, Ivet Bahar

## Abstract

The biological effects of human missense variants have been studied experimentally for decades but predicting their effects in clinical molecular diagnostics remains challenging. Available computational tools are usually based on the analysis of sequence conservation and structural properties of the mutant protein. We recently introduced a new machine learning method that demonstrated for the first time the significance of protein dynamics in determining the pathogenicity of missense variants. Here we present a significant extension that integrates coevolutionary data from Pfam database and we also introduce a new interface (*Rhapsody*) that enables fully automated assessment of pathogenicity. Benchmarked against a dataset of about 20,000 annotated variants, the methodology is shown to outperform well-established and/or advanced prediction tools. We illustrate the utility of our approach by *in silico* saturation mutagenesis study of human H-Ras. The tool is made available both as a webtool (rhapsody.csb.pitt.edu) and an open source Python package (<u>pip install prody-rhapsody</u>).

## Introduction

Single Nucleotide Polymorphisms (SNPs) are single DNA base pair changes that are inherited (germline variants) or occur during the organism’s lifetime (somatic variants). A SNP located in a coding region of the DNA may lead to translation of the gene codon into a different amino acid than the wild-type (non-synonymous SNPs), giving rise to a Single Amino acid Variant (SAV or missense variant). Both synonymous and non-synonymous SNPs can perturb the normal activity of a cell. For example, synonymous SNPs, can affect splicing, regulatory mechanisms and gene and/or protein expression levels although they do not affect the gene product’s sequence. SAVs can additionally have molecular effects in multiple ways, e.g. by affecting a protein’s active sites, its interaction with other proteins, or its stability.

More than half of the mutations implicated in human inherited diseases are estimated to be associated with SAVs (Stenson et al., 2017). As a result, devising analytical and computational approaches for predicting the effect of point mutations has been of broad interest, but equally challenging due to complex interrelated effects in the cell. In recent years, it became evident that comprehensive approaches integrating multiple perspectives are the only viable solutions to achieve higher accuracy in pathogenicity predictions and to interpret experimental data at the molecular level. In the case of SAVs, this means understanding not only the significance of the mutated amino acid vis-à-vis the biochemical activity of the protein, often captured by sequence-based conservation models, but also its importance for the fold stability and conformational mechanics and interactions, both intra- and intermolecular (Ancien et al., 2018).

Sophisticated analytical tools that focus on protein sequence conservation and residue coevolution, such as context-dependent modeling of sequence evolution (Feinauer and Weigt, 2017; Hopf et al., 2017) reached a good level of maturity in recent years. These computational tools predict the pathogenicity of missense variants by estimating the mutant’s fitness based on the reconstructed evolutionary history of the protein and comparative studies of its orthologs.

In contrast, structure-based modeling approaches have been lagging behind, even though classifiers that take account of 3D structures have shown considerable success (Adzhubei et al., 2010; Ancien et al., 2018). This class of computations has been limited by two factors: first, they are possible only when the 3D structure of the protein is known, either from experiments or from comparative modeling. Second, even when a structure is available, the traditional methods to investigate the effect of missense variants require expensive computations such as molecular dynamics (MD) simulations.

Such simulations do not lend themselves to genome-scale analyses; they are applicable on a case-by-case basis only and are limited by the time and space limitations of MD simulations.

Yet, recent years have seen a rapid growth in the structural characterization of the proteome with advances in structure determination (e.g. cryo-EM) technologies. In parallel, computationally efficient methods such as those based on Elastic Network Models (ENMs) have been developed, which efficiently provide insights into the intrinsic dynamics of proteins uniquely defined by their inter-residue contact topology (Bahar et al., 2010). Many analytical tools have been developed within the framework of ENMs, which focus on different aspects of protein equilibrium dynamics, both on a local (e.g. fluctuations in residue positions) and a global (e.g. coupled domain movements and allosteric switches) scale. ENMs have been broadly used for mechanistic studies of protein activities, but their potential utility in genome-scale studies of the impact of mutations became clear only in recent studies (Ponzoni and Bahar, 2018; Rodrigues et al., 2018).

The rapidly growing experimental data on the functional impact of missense variants and on protein structures provide a unique opportunity for building upon that first generation of pathogenicity predictors to develop and make accessible a classifier trained not only on well-established sequence- and structure-dependent properties, but also on *intrinsic dynamics*, derived from ENMs. A first attempt in that direction (Ponzoni and Bahar, 2018) yielded promising results and paved the way to the current development and implementation of *Rhapsody*, an advanced tool and user-friendly server for *<u>R</u>*apid *<u>h</u>*igh-*<u>a</u>*ccuracy *<u>p</u>*rediction of *<u>S</u>*AV *<u>o</u>*utcome based on *<u>dy</u>*namics, accessible at rhapsody.csb.pitt.edu.

The inclusion of dynamics-based features distinguishes *Rhapsody* from tools broadly used in the field for evaluating the pathogenicity of SAVs, e.g. PolyPhen-2 (Adzhubei et al., 2010), SIFT (Ng and Henikoff, 2003), CADD (Kircher et al., 2014) and others [see (Grimm et al., 2015) for a comparative review]. In addition to sequence-, structure- and dynamics-based features (Ponzoni and Bahar, 2018), the underlying algorithm now (i) incorporates conservation and *coevolution* features extracted from Pfam domains, inspired by the success of recent studies in the field (Feinauer and Weigt, 2017; Hopf et al., 2017), (ii) utilizes a refined dataset of about 20,000 human missense variants, built from consensus between clinical interpretations of variants found in multiple databases (DBs), (iii) is implemented as a standalone package, which may be used in conjunction with our *ProDy* API (Bakan et al., 2014, 2011). The server offers the option of using as input customized PDB structures, such as those stabilized under different conformational and oligomerization states along an allosteric cycle as well as those resolved for orthologues or obtained by comparative modeling.

The utility of *Rhapsody* is illustrated by way of application of human H-Ras, a highly conserved G-protein belonging to Ras subfamily of small GTPases, and comparison with deep mutational scanning data for this protein reported by Kuriyan, Ranganathan and coworkers (Bandaru et al., 2017). The new tool provides not only an efficient independent assessment of the pathogenicity of mutations, but also mechanistic insights into the molecular basis of the observed and/or predicted effect.

## Results

### Development of a new generation of dynamics-based pathogenicity predictors

We consider three groups of features, sequence-based (SEQ), structure-based (STR) and dynamics-based (DYN), in a Random Forest classifier (Ponzoni and Bahar, 2018). In the original version of the algorithm (“reduced” classifier in **Fig 1A**), SEQ features were computed by the PolyPhen-2 server (Adzhubei et al., 2010), STR features by using structural data from the Protein Data Bank (PDB) (Berman et al., 2000) and DYN features by the *ProDy* API (Bakan et al., 2014, 2011). This classifier was shown in previous work to achieve accuracy levels comparable to, if not better, than 11 existing tools (Ponzoni and Bahar, 2018). The current “full” version also evaluates the site entropy and coevolution properties of the mutated amino acid using Pfam domains (El-Gebali et al., 2019) and uses scores from BLOSUM62 amino acid substitution matrix (Henikoff and Henikoff, 1992). A detailed description of the features is presented in **Table S1** and in the Methods section. As will be presented below in more details, the “full classifier” outperforms the “reduced” classifier, based on various metrics (**Fig 1B**). This comes with a cost, however: the coverage is reduced as the tool requires that a variant be localized in a conserved Pfam domain of a protein.

**Fig 1:**
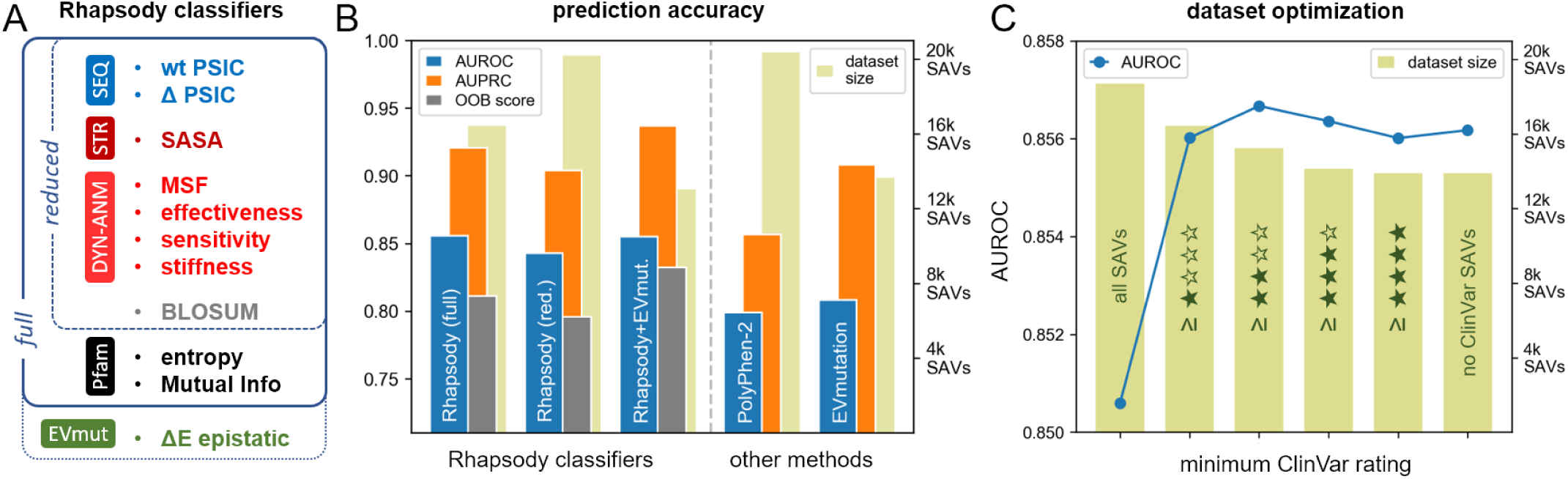
Rhapsody features and prediction accuracy. (A) List of random forest features organized into different classification schemes. See Methods and **Table S1** for detailed descriptions. (B) Accuracy measurements of the three *Rhapsody* classification schemes compared to two other representative prediction tools, PolyPhen-2 and EVmutation. *Light green bars* (*right ordinate*) in the background represent the relative size of the datasets of variants used for cross-validation, in the case of *Rhapsody*, and testing for the other two methods. *Blue, orange* and *gray bars* represent the results from three metrics of accuracy, Area Under ROC curve, Area Under Precision-Recall Curve and Out-Of-Bag score, respectively (*left ordinate*). See Methods and **Table S2** for dataset composition. (C) Impact of excluding variants from the training dataset on the “full” *Rhapsody* classifier, based on their rating (review stars) in the ClinVar DB. The *bars* represent the numbers of SAVs (*right ordinate*) that could actually be processed by *Rhapsody*, in different subsets distinguished by different numbers of “stars” (or reliability levels) reported in ClinVar. The *leftmost* bar refers to the “complete” set with of 0 - 4 stars; the 2^nd^ bar excludes those with zero stars; the 3^rd^ excludes those with 0 and 1 stars, and so on. The *blue curve* (*left ordinate*) displays the corresponding prediction accuracy levels (*left ordinate*).

### Construction of an integrated dataset

A dataset of labelled (deleterious/neutral) human SAVs for training the algorithm has been generated by combining five publicly available datasets [HumVar (Adzhubei et al., 2010), ExoVar (Li et al., 2013), PredictSNP (Bendl et al., 2014), VariBench (Thusberg et al., 2011), SwissVar (Mottaz et al., 2010)] with the Humsavar DB of all human missense variants annotated in the UniProtKB/Swiss-Prot DB (“UniProt: a worldwide hub of protein knowledge,” 2019) and the ClinVar archive of reports on the level of concordance between human variations and phenotypes (Landrum et al., 2016). **Table S2** provides information on the content of these datasets and their level of agreement. After filtering out discordant labels, we obtained a new “Integrated Dataset” (**IDS**) of 87,726 SAVs, of which 27,655 could be mapped onto PDB structures, a prerequisite for computing STR/DYN features, and 23,085 had PDB structures with at least 150 residues (see below for details about the choice of an optimal lower cutoff for the size of PDB structure).

The ClinVar DB provides a reliability assessment for each variant interpretation, on a scale between 0 to 4 (best) “review stars”, based on the number and consensus between the sources. In a preliminary analysis (**Fig. 1C**), we measured the average Area Under the ROC curve (AUROC) attained by the full classifier in a 10-fold cross-validation procedure while gradually excluding from the IDS those SAVs with lower rating. Notably, the exclusion of SAVs with 0 stars (assigned to variants with “no assertion” or “no assertion criteria provided”) resulted in a sizeable increase in accuracy (*blue curve* in **Fig. 1C**). This was followed by marginal improvements when excluding the 1-star (“single submitter” or “conflicting interpretations”) SAVs and a plateau/decrease after excluding the 2-star (“no conflicts and multiple submitters”), 3-star (“reviewed by experts”) and finally 4-star (“practice guideline”) SAVs. We therefore elected to exclude variants with 0-star ClinVar ranking only, which accounted for ∼12% of cases. The final, optimized dataset (**optIDS**) is thus composed of 20,361 SAVs with at least 1 ClinVar review star and 150+-residue PDB structures.

### Cross-validation and comparison with other tools

The optimized integrated dataset (optIDS) of 20,361 SAVs was used for evaluating the accuracy of the classifier through cross-validation. In **Fig. 1B**, we compare the performances of three variants of *Rhapsody* against two other methods, PolyPhen-2 (Adzhubei et al., 2010) and EVmutation (Hopf et al., 2017). The former is a broadly used tool for predicting the functional effects of human variants, which relies on a supervised naïve Bayes classifier trained on annotations, conservation scores and structural features that characterize the amino acid substitution. It is chosen here as a representative tool among several other publicly available tools [many of which we considered and compared in our earlier work (Ponzoni and Bahar, 2018)] because of its widespread use. The second method, EVmutation, is arguably the most accessible and powerful among the recent wave of tools that leverage coevolution analysis for predicting the fitness of mutants, going beyond the limitations of conservation analyses by taking account of the inter-dependencies between pairs of sequence positions. The change in “evolutionary statistical energy” ΔE incurred upon mutation is directly interpreted as a proxy for the mutant fitness. However, a cutoff energy for binary classification of mutants as deleterious or neutral is not defined.

The colored bars in **Fig. 1B** represent different metrics used for assessing the accuracy of each method. For the three *Rhapsody* variants on the left, we calculated the average AUROC, AUPRC (Area Under Precision-Recall Curve) and OOB (Out-Of-Bag) score from a 10-fold cross-validation on optIDS, while for PolyPhen-2 and EVmutation we plot the individual AUROC and AUPRC over the same dataset of variants. The *light green bars* in the background indicate the actual number of SAVs that could be evaluated by each approach. From the comparison, we notice that the “full” *Rhapsody* classifier outperforms both PolyPhen-2 and EVmutation according to both AUROC and AUPRC metrics. The introduction of Pfam-derived features, however, comes at the cost of a slightly decreased coverage, since Pfam domains often do not encompass the full span of a protein sequence, but only those portions that are preserved across species. In this regard, we notice that PolyPhen-2 has the widest scope, being able to return a prediction even for variants without a PDB structure.

Beside the full and reduced versions of *Rhapsody*, we also considered a third option, which incorporates the EVmutation “epistatic” score ΔE into the feature set (“*Rhapsody*+EVmut”). This variant slightly improved the Precision-Recall tradeoff and OOB score, but it also reduced the number of SAVs it could be applied to. Of note, the integration of EVmutation and *Rhapsody* leads to significantly more accurate predictions than EVmutation used alone.

### Contribution of selected features

**Fig. 2A** illustrates the relative weights of features in the “*Rhapsody*+EVmut” version of the classifier. In parallel with previous findings (Ponzoni and Bahar, 2018), sequence-based features (wtPSIC, ΔPSIC and entropy of Pfam domain) rank higher than dynamics-based (ENM-derived) features, since the latter lack residue specificity and cannot distinguish between different types of amino acid substitutions. Dynamics-based features, in turn, prove to be more informative than a widely used structural property, solvent accessibility (SASA). We note that these features are not necessarily independent. The heat map in **Fig. 2B** provides a quantitative description of their similarities. Yet, their explicit inclusion in the training algorithm has assisted in increasing prediction accuracies. We note in this context that a remarkable weight difference between two coevolution properties, the “ranked” Mutual Information and EVmutation’s ΔE score, is also evident. We emphasize that the former was chosen for its simplicity, which makes it orders of magnitude faster to evaluate computationally than EVmutation scores, for which a DB of precomputed values was used in practice (Hopf et al., 2017). For real-time evaluation of coevolution properties, the integration of more efficient coevolution algorithms might be envisioned.

**Fig 2:**
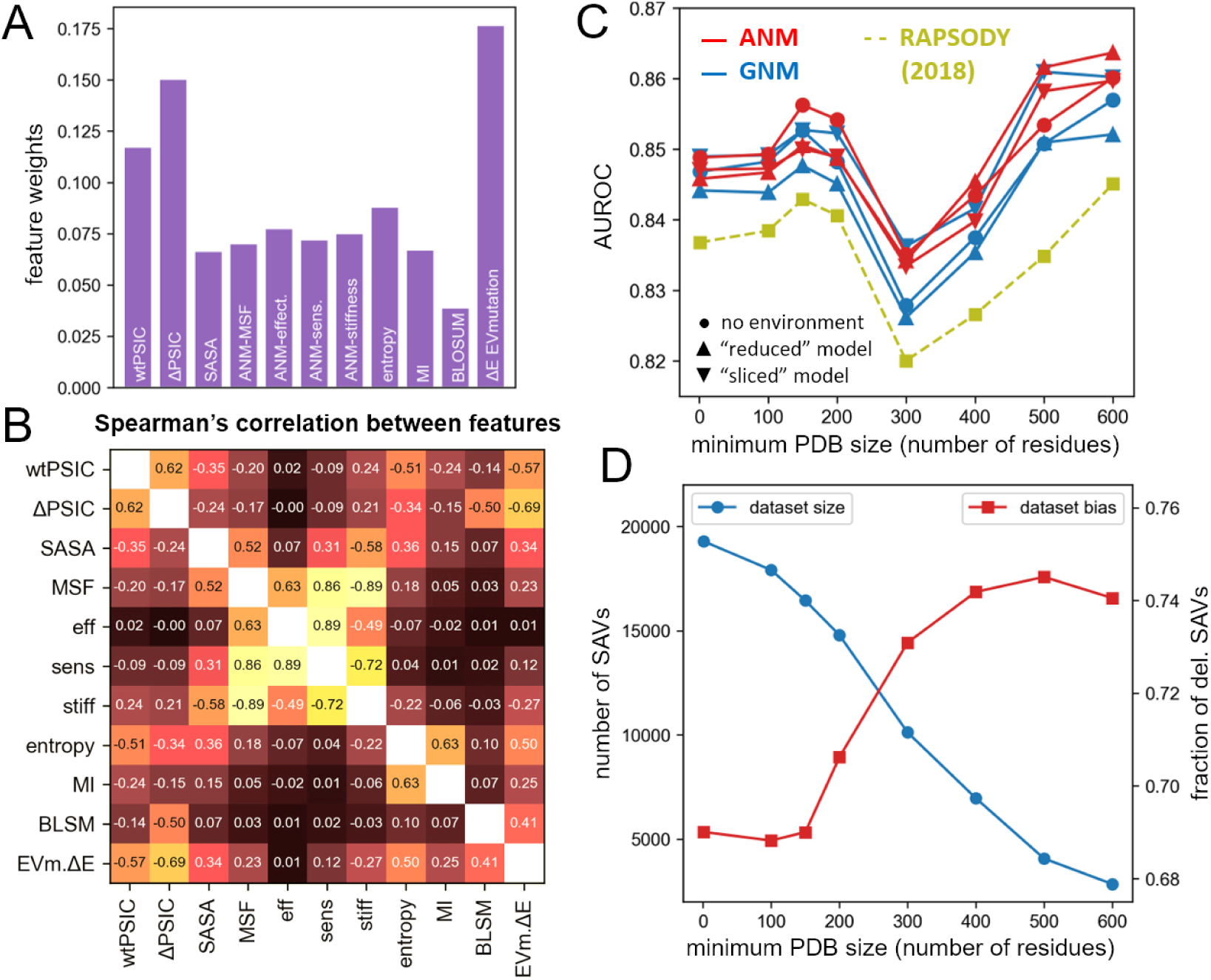
Analysis of the Rhapsody classifier. (A) Relative weights of features from the training of the *Rhapsody* classifier including EVmutation’s ΔE score. See also **Figure 2—figure supplement 1**. (B) Spearman’s correlations between all considered features. (C) Cross-validation of different versions of the “full” *Rhapsody* algorithm (GNM vs ANM dynamics features, with and without environmental effects), on different subsets of the Integrated Dataset obtained by setting a minimum PDB structure size (i.e. number of resolved residues). In *yellow*, we show the performance of the previous version of the algorithm (RAPSODY) (Ponzoni and Bahar, 2018). See also **Figure 2—figure supplement 2**. (D) Training dataset size (*blue*) and fraction of positive training examples (i.e. deleterious SAVs, in *red*) as a function of the minimum number of residues used to filter PDB structures based on their size.

### Higher accuracy achieved with larger structures

**Fig. 2C** illustrates the dependency of pathogenicity prediction accuracy on the minimum size of the PDB structure included in the evaluation of the STR and DYN features. An initial improvement in accuracy is observed when excluding structures with less than 150 residues and again when limiting the analysis to bigger structures with at least 500 residues. Such increased accuracy might be attributed to the importance of filtering out partially resolved protein structures and fragment that may not be suitable for computing overall structural and dynamical features. On the other side, the changes in the training dataset size and composition (*blue* and *red* curves, respectively, in **Fig. 2D**) might prevent a more direct detection of a correlation between PDB structures’ size and prediction accuracy. The non-monotonic behavior of the AUROC plot in **Fig. 2C** observed after the 150-residue cutoff could be attributed to the accentuated imbalance between deleterious and neutral variants in the training dataset. For these reasons, we deemed safe to set a minimum PDB size of 150 residues for our training examples (optIDS), where both dataset size and imbalance are maintained at more reasonable levels.

In **Fig. 2D**, we also note the increased accuracy achieved in the full version (set of *red/blue curves* for different ENM models, see Methods for details) compared to the reduced version (*yellow curve*), that resembles the original implementation of the algorithm (Ponzoni and Bahar, 2018) which did not include coevolutionary features.

Overall, these results confirm the usefulness of including intrinsic dynamics features in the context of functional assessment of variants, and further demonstrate the power of adopting an integrative approach that incorporates coevolution analysis into supervised learning approaches, thus taking advantage of its superior predictive power compared to single amino acid conservation properties.

## Application to H-Ras and comparison with experimental and clinical data

### Saturation mutagenesis analysis of human H-Ras protein

Kuriyan, Ranganathan and workers recently presented results from deep mutational scanning of human H-Ras (Bandaru et al., 2017), a highly conserved signaling protein which transduces signals through a nucleotide-dependent switch between active (GTP-bound) and inactive (GDP-bound) conformations. In order to recreate the natural, wild-type environment of this G protein as closely as possible, they isolated a “minimal biochemical network” comprised of one of the protein’s effectors (Raf) and two regulatory factors (GAP and GEF). The impact of a single mutation on the protein’s normal activity was then experimentally linked to the survival of the hosting bacterial system and quantified by a “fitness score” (ΔE). Different contexts have been considered and compared, by excluding one (“attenuated Ras”) or all (“unregulated Ras”) regulatory elements and by introducing a background oncogenic mutation (“Ras-G12V”). For the purpose of our analysis, we mainly focus on the complete (“regulated Ras”) experimental setup, designed to include those regulatory factors that might constrain Ras sequence variability and that are necessary to obtain a realistic assessment of mutants’ fitness.

**Fig. 3** presents the results from our so-called “*in silico* saturation mutagenesis” analysis. The results are presented in a 20 × *N* heatmap (**Fig. 3A**), where the entries are the color-coded pathogenicity probabilities (see **Figure 3—figure supplements 3**) predicted for all 19 possible substitutions at each of the *N* = 171 structurally resolved sequence positions of H-Ras (Uniprot sequence ID: P01112). The entries corresponding to the wild type amino acids are in *white*. The table structure mirrors that of analogous tables of experimental fitness measurements (Bandaru et al., 2017).

**Fig 3:**
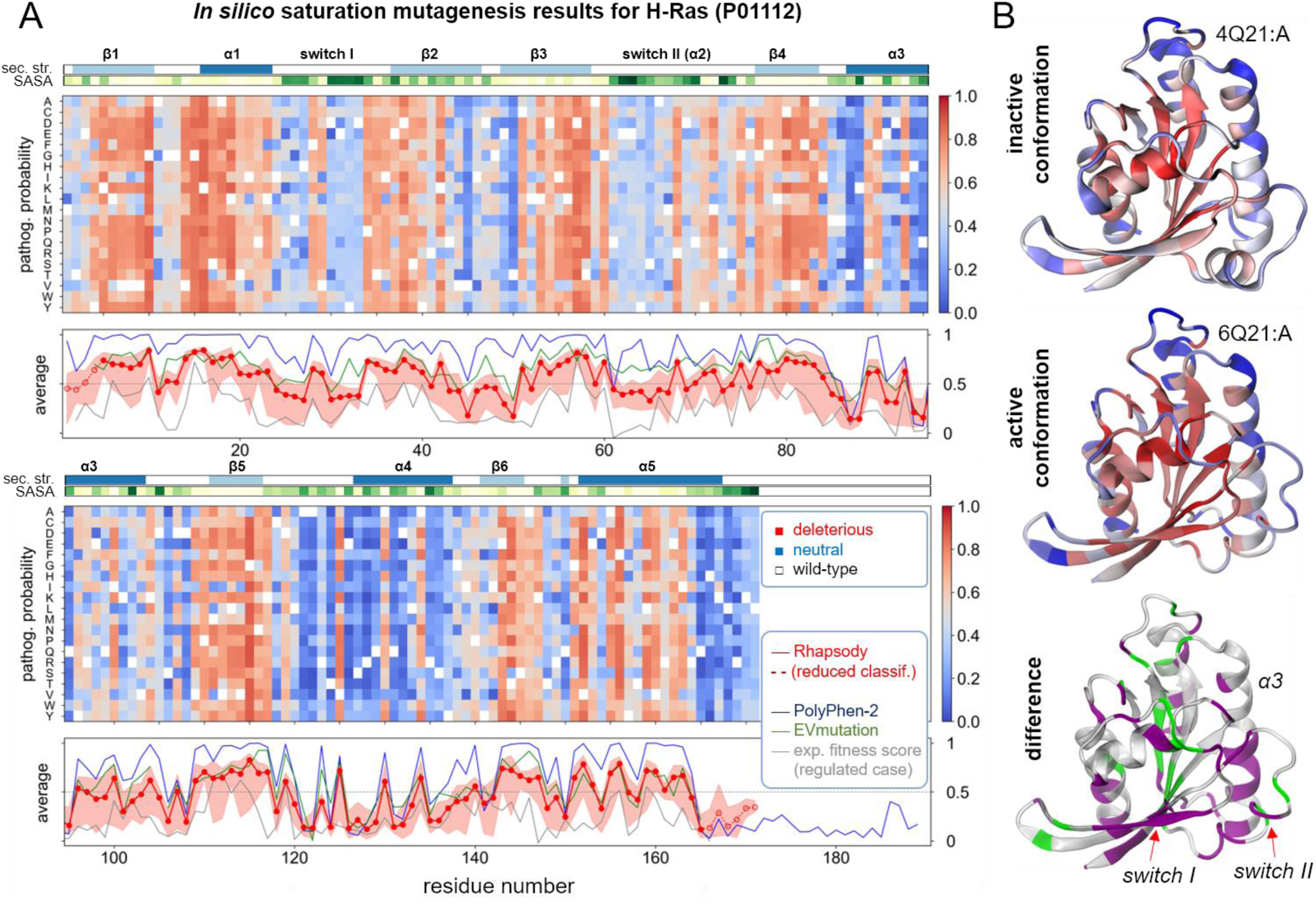
In silico saturation mutagenesis results for human H-Ras. (A) The predicted pathogenicity probabilities for all possible SAVs in H-Ras computed by *Rhapsody* are shown as a heatmap with a color code ranging from *red* (deleterious) to *blue* (neutral); see Methods for more details about the definition of pathogenicity probability. The corresponding residue-averaged pathogenicity profile is shown in *red* in the bottom panel, compared to analogous profiles from PolyPhen-2 (*blue*) and EVmutation (*green*) and from experimental fitness measures (*grey*). The two strips along the upper abscissa of the heatmaps display the secondary structure and solvent accessibility (SASA) along the sequence. The *Rhapsody* results are obtained for the structure in active state. The counterpart for the inactive form is presented in **Figure 3—figure supplement 1**. (B) Residue pathogenicities displayed by color-coded ribbon diagrams for active (*top panel*) and inactive *(middle panel*) H-Ras. *Red* and *blue* colors indicate the regions with high and low propensities for pathogenicity, respectively. The difference is shown in the *bottom* panel, with the respective *purple* and *green* regions referring to sites exhibiting increased and decreased pathogenicities in the active form. Note that the *purple* regions include the two switches involved in activation.

The structure-dependent (STR) and dynamics-based (DYN) features required by *Rhapsody*, were computed on the active, GTP-bound conformation of H-Ras (PDB ID: 6Q21, chain A). If allowed to automatically select a PDB structure, *Rhapsody* would select one of the inactive conformations (PDB ID: 4Q21, chain A), based on structure size alone. Repeating the computations for the latter showed that the predictions are very similar (see **Figure 3—figure supplements 1 and 2**), with the main differences localized at the switches I and II (**Fig. 3B**). These results are consistent with the robustness of ENM results to structural details, i.e. H-Ras structural dynamics is predominantly defined by its 3D fold, which defines its inter-residue contact topology, which in turn determines the intrinsically accessible spectrum of motions. Therefore, the potential impact of SAVs on collective mechanics can be inferred from either active or inactive form, provided that they retain the same overall fold.

At first glance, the heat maps in **Fig. 3A** show an alternating pattern of *blue* (neutral) and *red* (pathogenic) vertical bands that loosely correlate with either secondary structure or surface exposure of residues (*top strips*). Such a pattern can also be discerned in the bottom panels of **Fig. 3A**, which show the *residue-based pathogenicity profiles* (*red line*) computed upon averaging the entries in the corresponding column of the maps. Analogous profiles obtained using PolyPhen-2 (*blue*), EVmutation (*green*) and experimental fitness scores for “regulated-Ras” (Bandaru et al., 2017) (−ΔE, *gray*) reveal an overall agreement between computational predictions and experimental data (see also **Table S3**).

Comparison with experimental data shows that *Rhapsody* performs slightly better than EVmutation and PolyPhen-2 on the residue-averaged pathogenicity predictions. The Spearman’s rank-order correlations between computational results and experimental data for the “regulated” case (**Fig. 4A**) are |ρ| = 0.60 and 0.57 for *Rhapsody* predictions based on the inactive and active states, respectively, as opposed to |ρ| = 0.52 and 0.51 with EVmutation and PolyPhen-2; and both *Rhapsody* and EVmutation perform better than PolyPhen-2 in predicting individual fitness scores (|ρ| ≈ 0.42 *vs* 0.36). We also estimated the prediction accuracies using standard metrics such as AUROC and AUPRC. These, however, require a binary labelling of variants (neutral/pathogenic) that cannot be readily deduced from the distribution of experimental ΔE values, see **Figure 4—figure supplement 2**. We arbitrarily set the median of the distribution as a cutoff, while the 40^th^ and 60^th^ percentiles have been used to compute an uncertainty interval. The resulting ROC curves (**Fig. 4B**) confirm similar accuracy levels for *Rhapsody* and EVmutation, with respect to both individual (AUC) and residue-averaged (‹AUC›_res_) experimental data, and slightly lower accuracies for PolyPhen-2. Analogous conclusions emerge from the analysis of Precision-Recall curves, presented in **Fig. 5A and Figure 5—figure supplement 1**.

**Fig 4:**
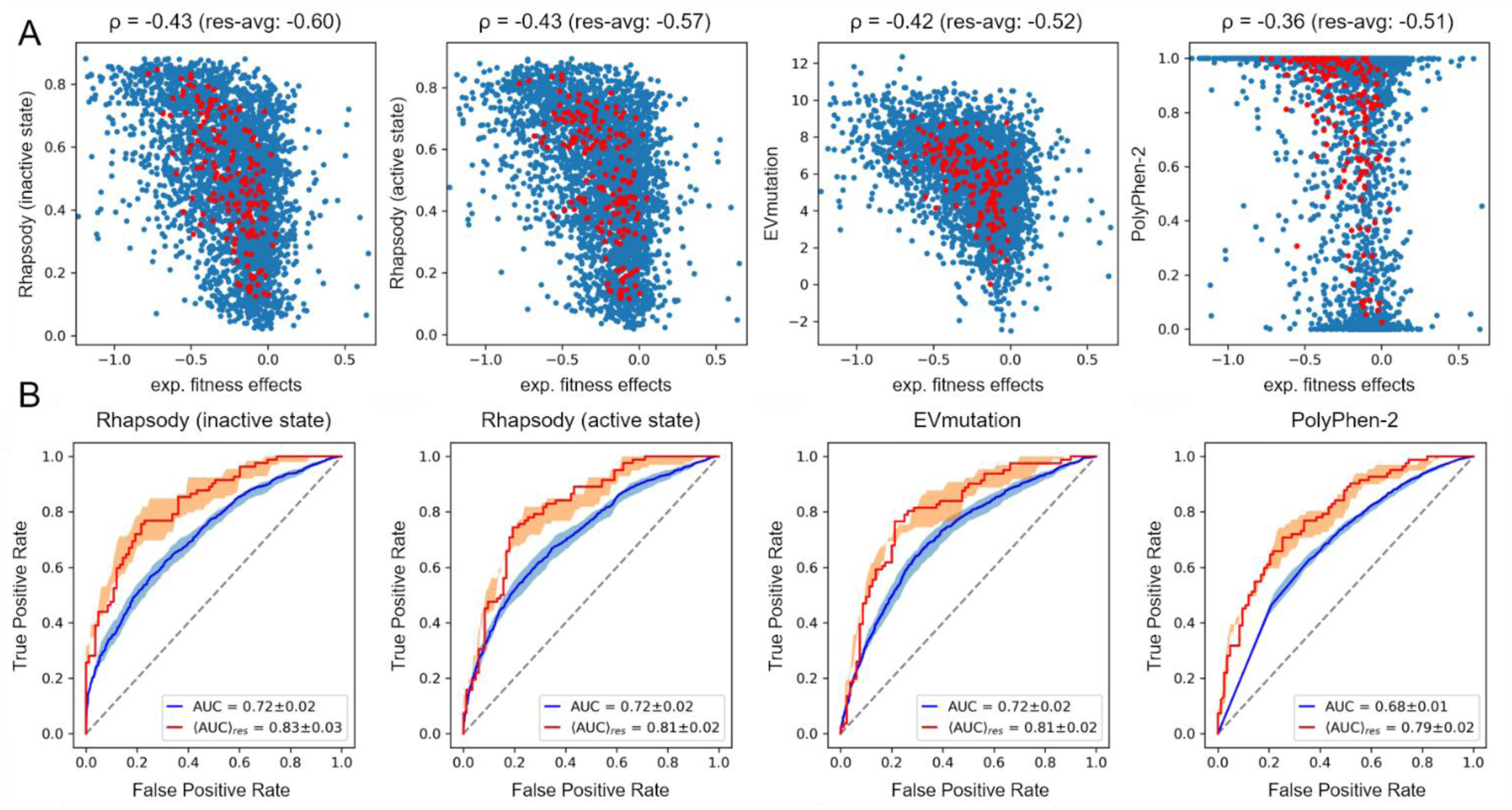
Pathogenicity predictions of human Ras protein variants. (A) Scatter plots and Spearman’s ρ correlations between experimental fitness scores from (Bandaru et al., 2017) and predictions from *Rhapsody* (based on inactive/active conformations), EVmutation and PolyPhen-2. *Red circles* correspond to residue-averaged values. See also **Figure 4—figure supplement 1**. (B) ROC curves for amino acid-specific (*blue*) and residue-averaged (*red*) predictions. The median of experimental ΔE measurements has been used as cutoff to assign binary labels to variants (see **Figure 4—figure supplement 2**). 40^th^ and 60^th^ percentiles have also been considered and used to compute uncertainty bands, represented in figure by semi-transparent *blue/red shades*. See also **Figure 4— figure supplement 3**.

**Fig 5:**
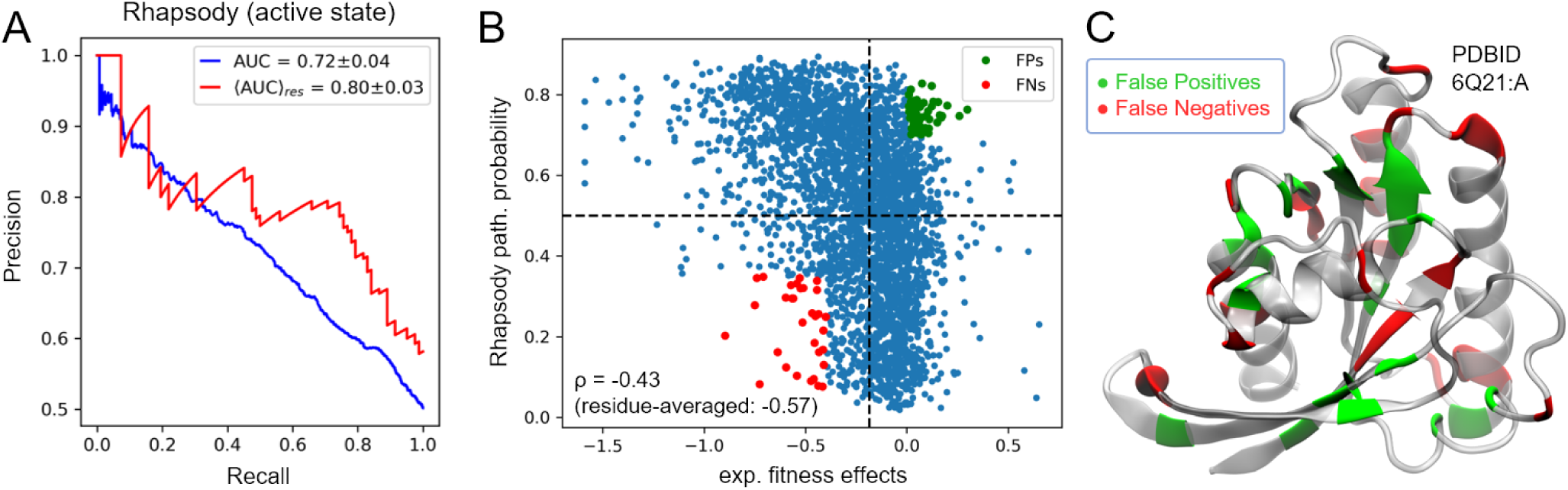
Analysis of Ras predictions. (A) Precision-recall plot for individual *(blue curve*) and residue-averaged (*red curve*) *Rhapsody* predictions of experimental fitness values. Corresponding AUCs are 0.72 and 0.80 respectively. Analogous plots for EVmutation and PolyPhen-2 are reported in **Figure 5—figure supplement 1.** (B) Scatter plot of *Rhapsody* predicted pathogenicity probabilities *vs* experimental measurements. (C) False Positives (*green*) and False Negatives (*red*) highlighted in plot (B) and displayed on the protein structure (active conformation).

Taken together, these results suggest that *Rhapsody* can be advantageously used for a first assessment of the regions that are most sensitive to mutations. Moreover, the consideration of a more diverse set of properties, such as dynamics-based features on top of sequence- and structure-based ones, as in *Rhapsody*, leads to improved prediction accuracy, and provides the opportunity of interpreting the observations in the light of structural and dynamic features of the investigated protein.

A visualization of *Rhapsody* incorrect predictions on Ras 3D structure (**Fig. 5B-C**) reveals that most False Negatives are localized on the protein’s surface, while False Positives are generally found in less exposed positions. A possible explanation is that the method is inherently biased towards the identification of residues important for the fold stability or internal dynamics, while locations subjected to other kinds of constraints, e.g. allostery and interactions with other proteins and small molecules, are more difficult to evaluate with the current set of features.

The systematic mischaracterization of gain-of-function variants (i.e. with ΔE > 0) in the “attenuated-Ras” and “unregulated-Ras” assays in **Figure 4—figure supplement 1**, prompted us to consider an alternative classification of variants’ effects as pathogenic or neutral, used for AUROC and AUPRC calculations. In principle, any deviation from wild-type activity levels induced by a variant might be viewed as a negative outcome—including an increased activity. We, therefore, considered an alternative labelling scheme where both significantly negative (loss-of-function) and positive (gain-of-function) fitness scores were interpreted as pathogenic, see **Figure 4—figure supplement 4**. Although performances of the classifiers do not change in most cases, we notice slightly higher accuracies in the “attenuated” cases (**Figure 4—figure supplement 5**). These results suggest that the interpretation of experimental data is of primary importance, and criteria for meaningful functional assessment of mutants must be set *a priori* based on the specific biological context.

### Analysis of H-Ras variants in gnomAD

We tested our predictions on a set of human variants found in healthy individuals, as collected by the gnomAD database (Karczewski et al., 2019). The assumption is that those substitutions seen in the 140,000 people tested (mostly normal population) are somewhat permissive. We therefore compared the distribution of predictions obtained by *Rhapsody* on this set of gnomAD SAVs with the corresponding fitness scores from the experimental study considered above (Bandaru et al., 2017).

The results, illustrated in **Fig. 6**, show that the predictions for the gnomAD SAVs are skewed towards “neutral” classification in both distributions, with 49 out of 82 total variants classified as “neutral” or “probably neutral” by our algorithm. Of note, 3 out of 4 “high count” SAVs (i.e. seen in 10 or more people) are interpreted as non-pathogenic by *Rhapsody*, while 2 out of 4 SAVs have a fitness score ΔE, as measured in the saturation mutagenesis study, significantly lower than the wild-type amino acid (when choosing the median of all values as cutoff).

**Fig 6:**
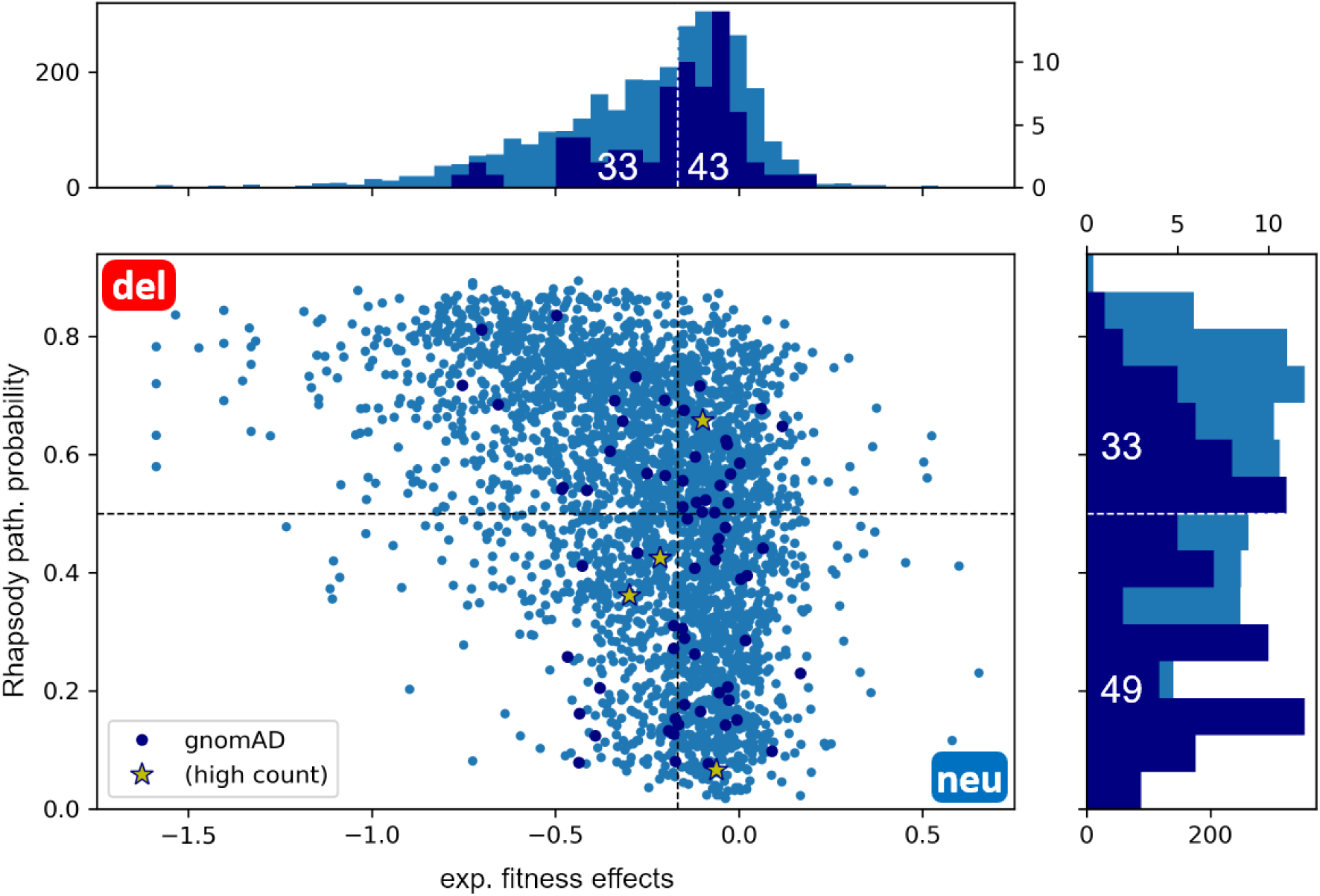
Analysis of H-Ras SAVs from gnomAD database. H-Ras SAVs (*dark blue dots*) collected from the gnomAD database found in healthy population are shown along with the results for all SAVS (*light blue dots*) on the scatter plot between *Rhapsody* pathogenicity probabilities and experimental fitness scores reported in (Bandaru et al., 2017). “High-count” SAVs (*yellow stars*) were seen in at least 10 individuals. The marginal plots show the corresponding distributions computed for all variants (*light blue*) and gnomAD variants (*dark blue*).

## Discussion

In this paper, we presented a novel machine learning approach for evaluating the functional impact, or “pathogenicity”, of human missense variants and illustrated its application to H-Ras. The newly introduced interface, *Rhapsody*, integrates dynamical features computed from ENMs of protein structures and attains a state-of-the-art prediction accuracy with a relatively simple design. We also highlighted how the method can be used not only for hypothesis generation (pathogenicity predictions of variants of unknown significance) but also for hypothesis testing, by providing a unified framework for comparing the predictive power of new as well as more established features. For instance, we demonstrated the potential of including in our machine-learning algorithm the ENM-derived dynamics-based features, in addition to more traditional conservation and structural features, and emphasized the need for a better integration with coevolution analysis, whose recent application to variant pathogenicity predictions produced outstanding results.

Through the analysis of saturation mutagenesis studies and other experimental and clinical data, we identified the strengths and limitations of our approach, and compared it against other prediction tools. We observed a general robustness of computational predictions, especially in the identification of residue sites that are sensitive to *any* mutation, regardless of the specific amino acid substitution. This information can be invaluable for the study of the functional mechanisms of proteins, especially when projected on the 3D structures. The use of structure-based properties, in combination with sequence conservation properties (“reduced” classifier), remains as a valid replacement for the more sophisticated coevolutionary analysis, whenever the latter cannot be applied due to lack of suitable multiple sequence alignments (MSAs).

The comparison with clinical and experimental data also revealed a few issues that, in our opinion, ought to be resolved in order to advance the field. Beside the more obvious shortcomings, such as the imbalance of available datasets towards pathogenic variants and the often-contradictory clinical interpretations found in different DBs, we reported our difficulties in interpreting data from more recent large-scale experimental studies. These studies provide a unique opportunity for dramatically increasing the size of training datasets. However, there is a need for a systematic definition of what is considered as a “pathogenic” variant, that would account for both loss-of-function and gain-of-function effects in relation to the biological context of the affected protein.

We expect future improvements to our method to address some of these shortcomings. Future developments will primarily focus on the definition of better structure-based features, for instance by using residue-specific ENMs (Kaynak et al., 2018), and on the identification and selection of more *context-dependent* structural data, e.g. the biological assembly rather than the asymmetric unit usually reported in the PDB. A recent ENM study has demonstrated how the consideration of the intact structures of multimers, complexes or assemblies improves the accuracy of predicted fluctuation spectrum of residues, and predictions from that server (*DynOmics*) (Li et al., 2017) could be used for evaluating context-dependent structural and dynamic properties. For example, a region that is deemed to be tolerant to mutations by virtue of its solvent-exposure in the PDB resolved structure, may actually represents a buried site in the physiologically relevant complex/assembly in which it participates, and a substitution at that region could alter its binding properties or interfacial packing. Another important property is protein-protein interactions. A recent study has demonstrated how disease-associated SAVs are likely to be located at singlet hot spots at protein-protein interfaces (Ozdemir et al., 2018). Consideration of the involvement of residues in interfacial interactions is expected to improve the prediction accuracy of current algorithms

We will also consider permanently including “coevolution-based” features into our algorithm, by either querying DBs of precomputed scores, as done with EVmutation, or by implementing interfaces to external libraries for Direct Coupling Analysis (DCA) [EVcouplings (Hopf et al., 2018), PconsC4 (Michel et al., 2018), Gremlin (Ovchinnikov et al., 2014), plmDCA (Ekeberg et al., 2014)]. A direct implementation would allow for much more flexibility in both fine-tuning the feature definitions, for instance by introducing normalizations that were shown to improve their predictive power (Feinauer and Weigt, 2017), and in the selection of MSAs for DCA calculations. MSAs could be either obtained from up-to-date DBs, such as Pfam, or built from scratch (as done by PolyPhen-2) whenever Pfam domains cannot be found, or even manually provided by the user. This upgrade would also make it possible to derive approximate dynamics information from the inferred contact map, thus circumventing the need for 3D structures (Butler et al., 2018), and to extend the analysis to non-human proteins.

Another possible improvement would be the consideration of the signature dynamics of the protein family to which the investigated protein belongs, as opposed to the dynamics of the protein alone (Zhang et al., 2019). In the same way as variations in sequence among family members point to sites that can, or cannot, tolerate mutations, family-based analyses can provide deeper insights into sites whose mechanistic properties are indispensable for function or for differentiation among subfamily members. Finally a decomposition of the mode spectrum and focus on the global modes could help extract information on high-energy localization (hot) spots, emerging as peaks in high frequency modes, which would likely be pathogenic if substituted by a larger size amino acid, as well as those residues at the flexible (hinge) region between domains, where an opposite effect (replacement of a small amino acid, like Gly or Ala, by a larger/stiffer one) may be detrimental (Dorantes-Gilardi et al., 2018; Rodrigues et al., 2018; Sayilgan et al., 2019). It remains to be seen if incorporation of such features in our random forest algorithm can assist in further improving the accuracy of pathogenicity predictions.

The *Rhapsody* algorithm is provided both as an open-source Python package (`pip install prody-rhapsody`) and a web tool (`rhapsody.csb.pitt.edu`). The latter has been designed as a user-friendly service that requires minimal user input or computing skills, but also allows for some customization, such as selecting or uploading a specific PDB structure. The *Rhapsody* webserver can be used for both obtaining predictions on a list of human missense variants (batch query) and for visualizing a complete *in silico* saturation mutagenesis analysis of a human sequence, akin to those presented in **Fig. 3** for H-Ras. Finally, the site offers tutorials, training data (optIDS) and precomputed features needed for reproducing all results presented here, or for analyzing new variants. The documentation also explains how to train a model on a completely different set of features and using a different training dataset, thus providing researchers with a flexible tool for analyzing personalized datasets and testing new predictors with the help of all the functionalities implemented in *Rhapsody*.

## Materials and Methods

### Datasets of SAVs

The detailed composition of the Integrated Dataset (IDS) of annotated SAVs used for training and testing of the *Rhapsody* classifier is shown in **Table S2**. Five datasets, namely HumVar (Adzhubei et al., 2010), ExoVar (Li et al., 2013), predictSNP (Bendl et al., 2014), VariBench (Sasidharan Nair and Vihinen, 2013) and SwissVar (Mottaz et al., 2010), were already considered in the first implementation of the algorithm (Ponzoni and Bahar, 2018) and the actual data and labels were extracted from a published review of prediction tools (Grimm et al., 2015). In addition, we included data from Uniprot’s Humsavar DB of annotated human missense variants (release: 2018_03) (“UniProt: a worldwide hub of protein knowledge,” 2019) and from ClinVar, a public archive of associations between human variants and phenotypes (release: Feb. 20, 2019) (Landrum et al., 2016). The resulting dataset, after discarding SAVs with conflicting interpretations, contained 87,726 SAVs, of which 27,655 could be mapped to a PDB structure, see **Table S2**. About 70% of SAVs were labelled as “deleterious”.

### Random forest and features

All features used by the random forest classifier are summarized in **Fig. 1A** and detailed in **Table S1**. In brief, the reduced classifier is similar to the original version presented in (Ponzoni and Bahar, 2018): it uses a combination of two sequence-based conservation features, extracted from the output of PolyPhen-2 (Adzhubei et al., 2010); one structural property, the solvent-accessible surface area computed by the DSSP algorithm (Touw et al., 2015) on PDB structures; four dynamical features [mean square fluctuation, perturbation-response scanning (Atilgan and Atilgan, 2009) effectiveness/sensitivity and mechanical stiffness (Eyal and Bahar, 2008) of a residue], based on ENM representation of protein structures [either Gaussian Network Model (GNM) (Li et al., 2016) or anisotropic network model (ANM) (Eyal et al., 2015)]; and substitution scores from the BLOSUM62 matrix (Henikoff and Henikoff, 1992). In addition to these, the full classifier also includes two sequence-based features computed on Pfam domains (El-Gebali et al., 2019): the Shannon entropy at the mutation site and its (ranked) average Mutual Information (MI) with respect to all other sites in the conserved domain. The latter incorporates coevolution effects, yet such a simplistic (and computationally feasible) approach does not significantly improve on single-site conservation analysis, see **Fig. 2A**. For this reason, we also considered a more advanced coevolution measure, the change ΔE in evolutionary statistical energy predicted by EVmutation (Hopf et al., 2017) epistatic model. Precomputed predictions of mutation effects provided by EVmutation have been both used for comparison and included as an additional feature in an expanded version of the classifier, see e.g. **Fig. 1B**.

When computing ENM features, the presence of other chains in the PDB structure can be accounted for either by including environmental effects in an “effective” way, i.e. by modifying the Hessian matrix [“reduced” model (Ming and Wall, 2005; Zheng and Brooks, 2005)], or by explicitly extending the system to include all chains in the complex and using the portion of eigenvectors corresponding to the investigated system (“sliced” model). We evaluated the performances of each model (see e.g. **Fig. 2C**, where different classification schemes are compared, while simultaneously increasing the minimum size of PDB structures): The results proved to be robustly maintained, irrespective of the selected model, apart from a slightly better accuracy using ANM. However, no effort has been made for selecting biological assemblies (e.g. actual functional complexes rather than, for instance, crystallographic asymmetric units), a further development that we deem worthy of future investigations.

The final score returned by the random forest classifier is converted to a “pathogenicity probability”, computed as the fraction of deleterious training examples that have been assigned with a similar score during cross-validation (see **Figure 3—figure supplement 3**). A pathogenicity class is also assigned based on a score cutoff chosen as to maximize the Youden’s index in the relative ROC curve computed during the cross-validation procedure. A similar criterion is used to define a binary classification from EVmutation’s epistatic score ΔE.

The two main hyperparameters of the random forest classifier, the number of trees and the max number of features considered when looking for the best split, were optimized through 10-fold cross-validation, see **Figure 2—figure supplement 3**. The open-source Python library *scikit-learn* was used for the implementation of the random forest classifier and accuracy metrics calculations.

## Data availability

The “Tutorials” section on the *Rhapsody* website (rhapsody.csb.pitt.edu) contains all the data and Python scripts needed to replicate the results illustrated in this paper. The training dataset (IDS) is also included in the public release of the *Rhapsody* package available on the Python Package Index (PyPI) and relative git repository (see “Download” section on the website) and can be used for the training of a customized version of the classifier. Step-by-step instructions for local installation and usage, and on how to expand and contribute to the *Rhapsody* package can be found in the “Tutorials” section as well.

## Acknowledgments

Support from NIH grants P41 GM103712 and P30 DA035778 is gratefully acknowledged by IB.

## Author contributions

The project was designed by LP and IB. The *in silico* models were generated by LP with input from IB. The *Rhapsody* Python package and webserver were implemented by LP. Clinical interpretations of the variants were done by ZNO. The manuscript was written by LP, ZNO and IB. All authors approved the manuscript.

