## Supplementary Information for "Rhapsody: Pathogenicity prediction of human missense variants based on protein sequence, structure and dynamics"

### Supplementary Tables

#### Table S1: Definition of features and their interpretation

<table>
<tr><th colspan="2">Classifier</th><th>Feature</th><th>Category</th><th>Definition</th><th>Interpretation</th></tr>
<tr><td rowspan="10">Full classifier</td><td rowspan="8">Reduced classifier</td><td>wild-type PSIC</td><td rowspan="2">sequence-based, computed by PolyPhen-2 (Adzhubei et al., 2010)</td><td>conservation of wild-type amino acid, based on the Position-Specific Independent Counts (PSIC) score (Sunyaev et al., 1999), which takes into account the background probability of observing a given amino acid at any position</td><td rowspan="2">conserved residues are more sensitive to mutations but sometimes can be replaced by equally conserved amino acid (Δ PSIC ≈ 0)</td></tr>
<tr><td>Δ PSIC</td><td>difference between PSIC scores of wild-type and mutated amino acids</td></tr>
<tr><td>SASA</td><td>Structure-based, computed by DSSP (Touw et al., 2015)</td><td>Solvent-Accessible Surface Area</td><td>buried residues are usually important for fold stability</td></tr>
<tr><td>MSF</td><td rowspan="4">Dynamics-based, computed by ProDy (Bakan et al., 2011) using ENMs</td><td>mean square fluctuations or displacements of $C_\alpha$'s calculated analytically from the normal modes of an elastic network model (ANM or GNM)</td><td>“hinge” (i.e. less mobile) regions play an important role in defining the mechanical aspects of a protein’s function</td></tr>
<tr><td>effectiveness</td><td rowspan="2">allosteric properties, computed with the Perturbation Response Scanning (PRS) (Atilgan and Atilgan, 2009) algorithm, related to the ability of a residue to transmit (sense) mechanical deformation signals to (from) distal sites</td><td rowspan="2">residues with high effectiveness (effectors) and either low or high sensitivity are generally more probable to be affected by mutations</td></tr>
<tr><td>sensitivity</td></tr>
<tr><td>stiffness</td><td>a measure of a residue’s resistance to uniaxial tension. Lower if the tension is accommodated upon reconfiguration along soft (low frequency) modes (Eyal and Bahar, 2008)</td><td>higher stiffness correlates with higher pathogenicity if mutated</td></tr>
<tr><td>BLOSUM</td><td>sequence-based</td><td>amino acid substitution score derived from BLOSUM62 (Henikoff and Henikoff, 1992)</td><td>more negative values correspond to lower probability for a given amino acid to replace the wild-type one without inducing negative effects</td></tr>
<tr><td rowspan="2"></td><td>residue entropy</td><td rowspan="2">sequence-based, computed by ProDy (Bakan et al., 2011) on Pfam domains (El-Gebali et al., 2019)</td><td>average Shannon entropy of the mutation site based on Multiple Sequence Alignments downloaded from Pfam database</td><td>lower entropy, typical of conserved residues, correlates with more sensitivity to mutations</td></tr>
<tr><td>ranked Mutual Information</td><td>ranking, normalized between 0 to 1, of the average mutual information between a given residue and all other Pfam domain sites</td><td>top ranking (rank ≈ 0) values correspond to residues with strong evolutionary co-dependency with other residues in the conserved domain</td></tr>
<tr><td colspan="2"></td><td>ΔE epistatic score</td><td>sequence-based, computed by EVmutation (Hopf et al., 2017)</td><td>change in evolutionary statistical energy that incorporates effects of pairwise epistasis (residue covariation)</td><td>negative ΔE values correspond to less probable mutant sequences (putatively deleterious) compared to wild-type sequence</td></tr>
</table>

**Table S2: Integrated Dataset composition and extent of overlap between different datasets (*)**

| | ClinVar | ExoVar | Humsavar | HumVar | PredictSNP | SwissVar | VariBench |
|---|---|---|---|---|---|---|---|
| **ClinVar** | **20,814** (89) **61.4%** | | | | | | |
| **ExoVar** | 2,984 (475) | **8,809** **58.5%** | | | | | |
| **Humsavar** | 19,043 (2,729) | 5,098 (341) | **68,386** **42.4%** | | | | |
| **HumVar** | 12,429 (2222) | 5,438 (21) | 37,003 (1005) | **40,177** **52.3%** | | | |
| **PredictSNP** | 299 (21) | 8 | 3,222 (48) | 25 (1) | **10,459** (1) **62.6%** | | |
| **SwissVar** | 1,119 (127) | 10 | 7,058 (60) | 56 (1) | 21 (1) | **8,862** (1) **31.7%** | |
| **VariBench** | 665 (291) | 1 | 697 (66) | 14 | 2 | 4 | **10,237** **42.1%** |

**Integrated Dataset**

- total number of SAVs: **94,505**
- SAVs successfully submitted to PolyPhen-2: **91,697**
- SAVs with unambiguous clinical interpretation: **87,726**
- SAVs mapped to PDB: **27,655** (deleterious SAVs' percent: **70%**)

*(*) On the diagonal, we show in boldface the size (number of SAVs) of each dataset, in parentheses the number of "ambiguous" (i.e. without a clear clinical interpretation) SAVs if any, and in red the dataset bias (percentage of deleterious SAVs). Off-diagonal cells contain the size of the intersection between the two datasets and in parentheses the number of SAVs with discordant interpretations. The inset shows information about the Integrated Dataset obtained by merging the 7 datasets.*

**Table S3: Spearman's correlations between experimental and computational pathogenicity/fitness scores observed or predicted for H-Ras (*)**

| | exp. fitness ("regulated") | Rhapsody (inactive) | Rhapsody (active) | EVmutation | PolyPhen-2 |
|---|---|---|---|---|---|
| **exp. fitness ("regulated")** | - | -0.426 | -0.428 | -0.42 | -0.364 |
| **Rhapsody (inactive)** | (-0.603) | - | 0.939 | 0.715 | 0.82 |
| **Rhapsody (active)** | (-0.570) | (-0.939) | - | 0.716 | 0.812 |
| **EVmutation** | (-0.518) | (-0.811) | (-0.788) | - | 0.703 |
| **PolyPhen-2** | (-0.513) | (-0.874) | (-0.862) | (-0.784) | - |

*(*) Values in parentheses in the lower triangular part of the table refer to residue-averaged quantities. Note that pathogenicity and fitness are inversely proportional, hence the negative sign in the entries corresponding to comparison of experimental and computational data.*

### Supplementary Figures

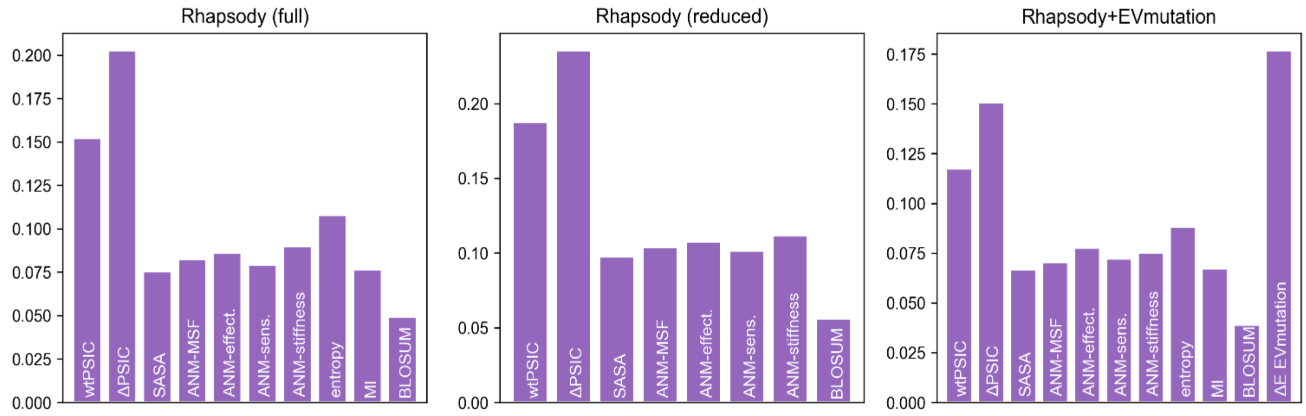


**Figure 2—figure supplement 1: Feature weights of the three Rhapsody classifiers**

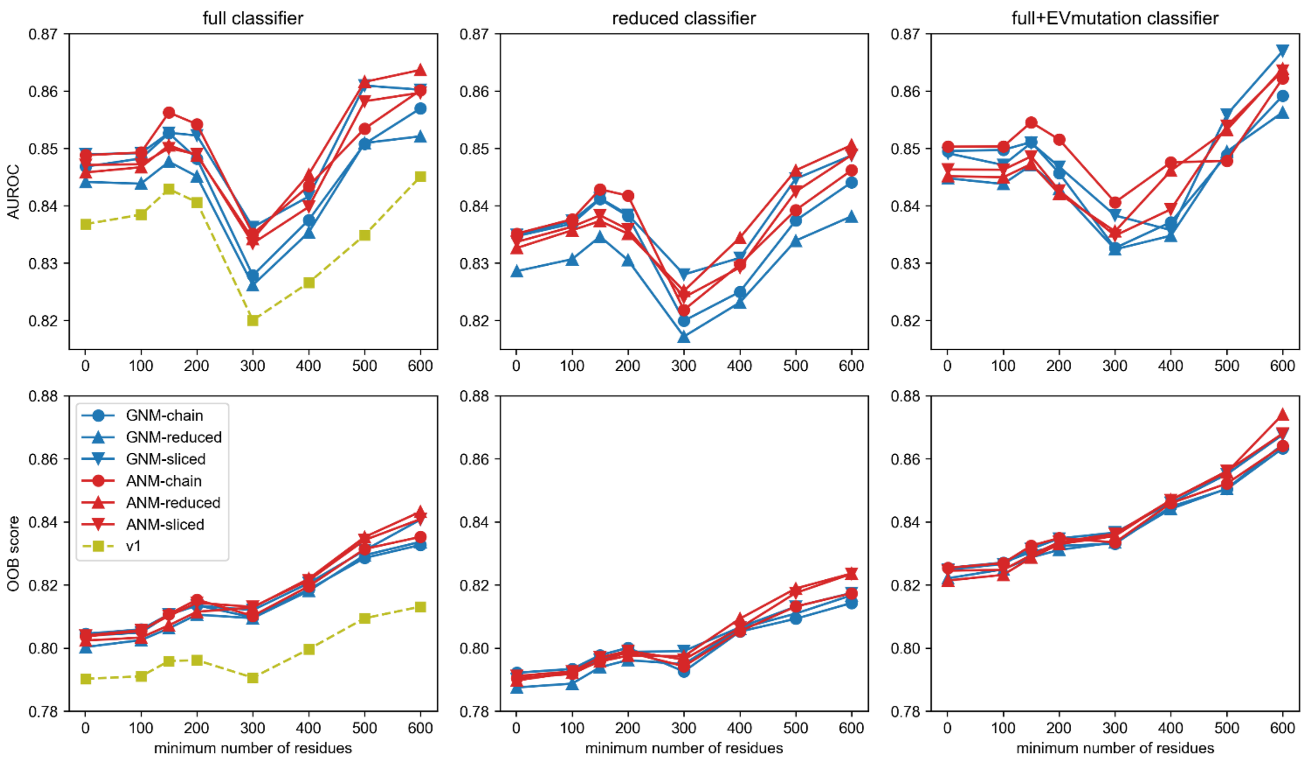


**Figure 2—figure supplement 2: Prediction accuracy *vs* minimum PDB structure size**

For each classifier ("full", "reduced" or with EVmutation score integrated) we tested different ways of computing dynamical features, based on GNM or ANM, and with (reduced/sliced models) or without (chain model) environmental effects. Each version has been tested through cross-validation on different subsets of the Integrated Dataset obtained by progressively increasing the minimum size requirement on PDB structures.

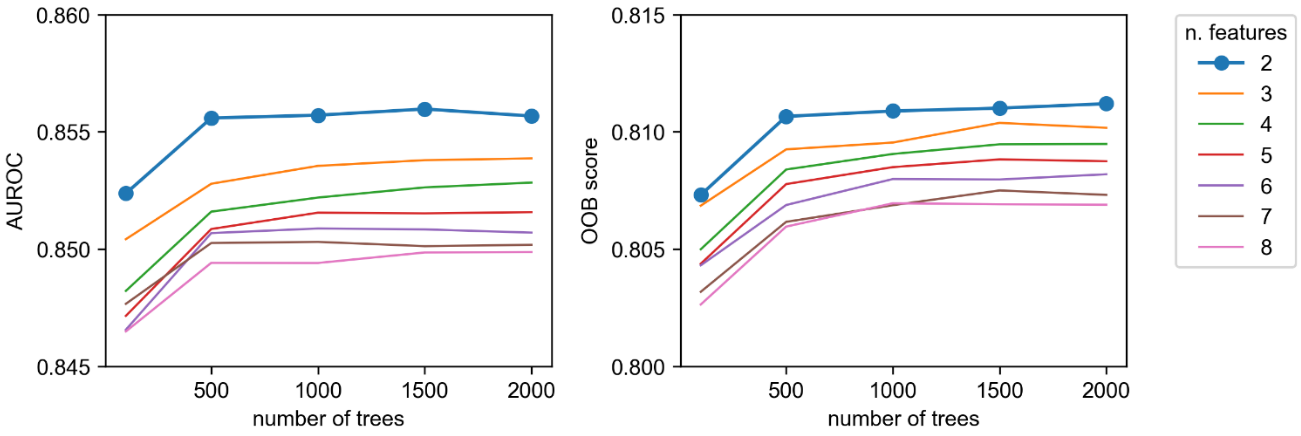


**Figure 2—figure supplement 3: Hyperparameter optimization**

The number of trees (or estimators) in the forest and the maximum number of features (considered for calculating the best split in each single decision tree) have been optimized through 10-fold cross-validation. Based on these measurements of AUROC and out-of-bag (OOB) score, we finally set those parameters to 1500 and 2, respectively.

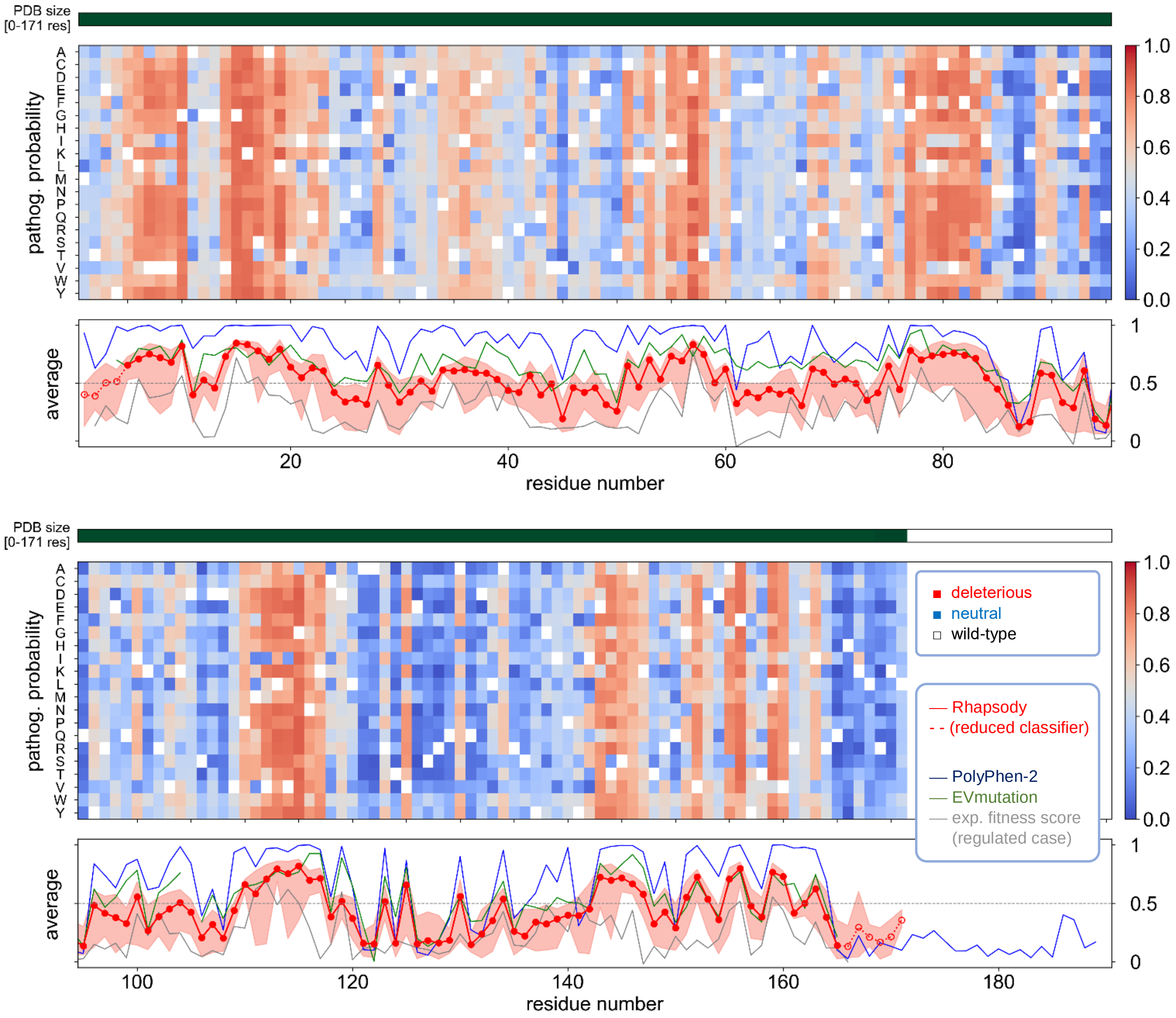


**Figure 3—figure supplement 1: *In silico* saturation mutagenesis analysis of human H-Ras protein in its inactive state**

When queried with a Uniprot accession number, Rhapsody automatically maps the sequence to one or more PDB structures, based on internal criteria (mainly, the structure size). In the case of human H-Ras (Uniprot ID: P01112), the sequence is mapped on a structure representing the GDP-complexed, inactive state (PDB ID: 4Q21, chain A) in 98% of residues (168 out of 171) and on another GDP-complexed structure (PDB ID: 1AA9, chain A) in the remaining residues. Almost all sequence could also be mapped on a single Pfam domain (PF00071), thus allowing the application of the full classifier (solid red line in the bottom panel).

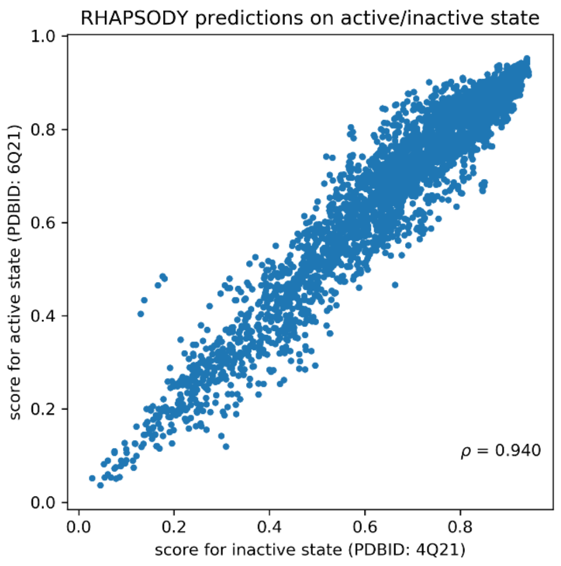


**Figure 3—figure supplement 2: H-Ras computational predictions based on active *vs* inactive conformations**

Comparison of the *Rhapsody* predictions (pathogenicity scores) computed by using the active (PDB ID: 6Q21) and inactive (PDB ID: 4Q21) conformations of H-Ras (Milburn et al., 1990) for evaluating the STR and DYN features used as input.

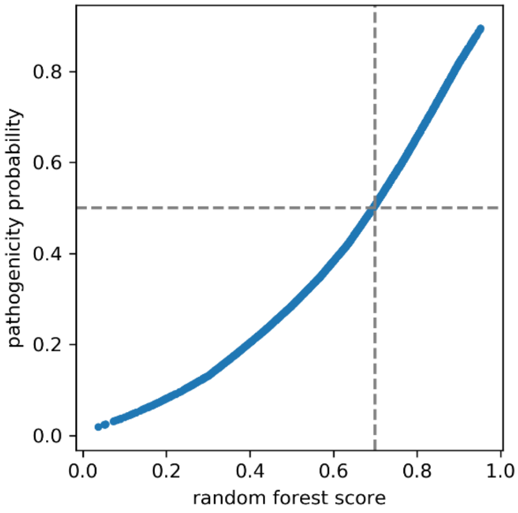


**Figure 3—figure supplement 3: Pathogenicity probability *vs* random forest score**

The score cutoff (≈0.7) used for assigning a pathogenicity class ("deleterious"/"neutral") is mapped to a cutoff around 0.5 in the pathogenicity probability.

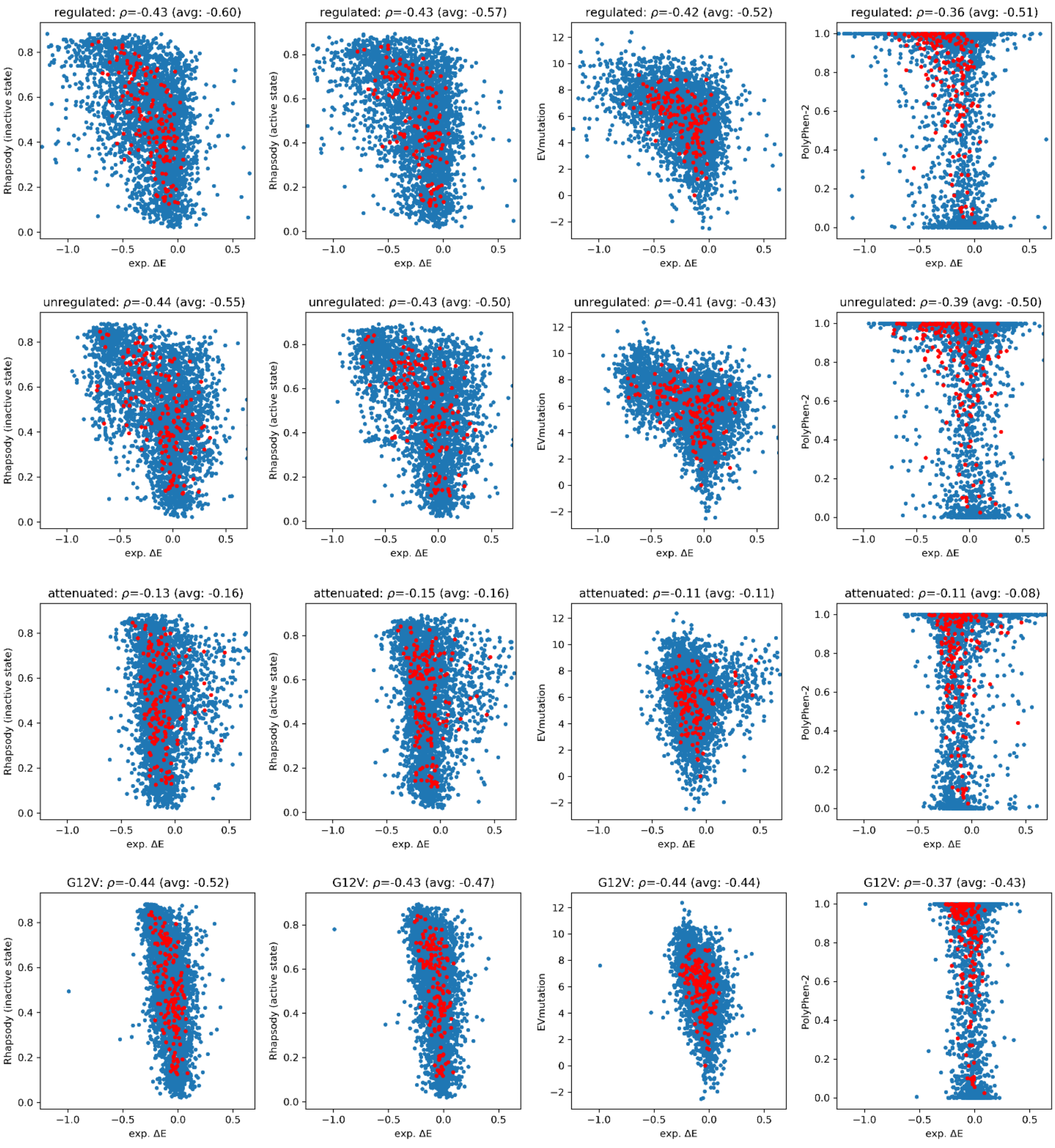


**Figure 4—figure supplement 1: Human H-Ras computational predictions *vs.* experimental fitness scores**

Extended version of **Fig. 4A** with all four experimental setups reported in (Bandaru et al., 2017). The first row is identical to **Fig. 4A** and is included here for completeness and comparison. The *blue dots* refer to the effect of specific substitutions of amino acids and *red dots* to the residue averages over all 19 substitutions.

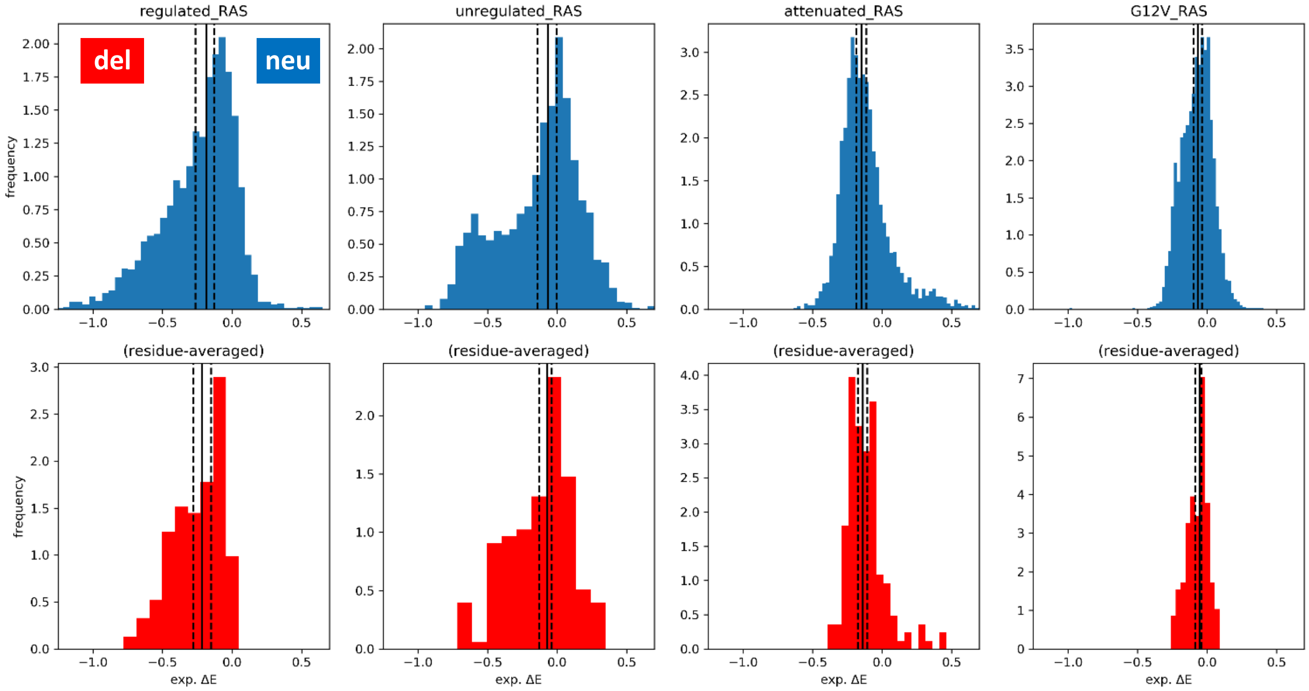


**Figure 4—figure supplement 2: Distributions of Ras experimental fitness scores.**

The experimental data collected under four different conditions (Bandaru et al., 2017) are used here to generate the histograms of fitness scores for all pairwise substitutions (*top* panels) and averages over all substitutions of a given residue (*bottom* panels). Solid and dashed lines mark the median and $40^{th}/60^{th}$ percentile values, respectively, used for classifying variants into pathogenic (*left*) and neutral (*right*).

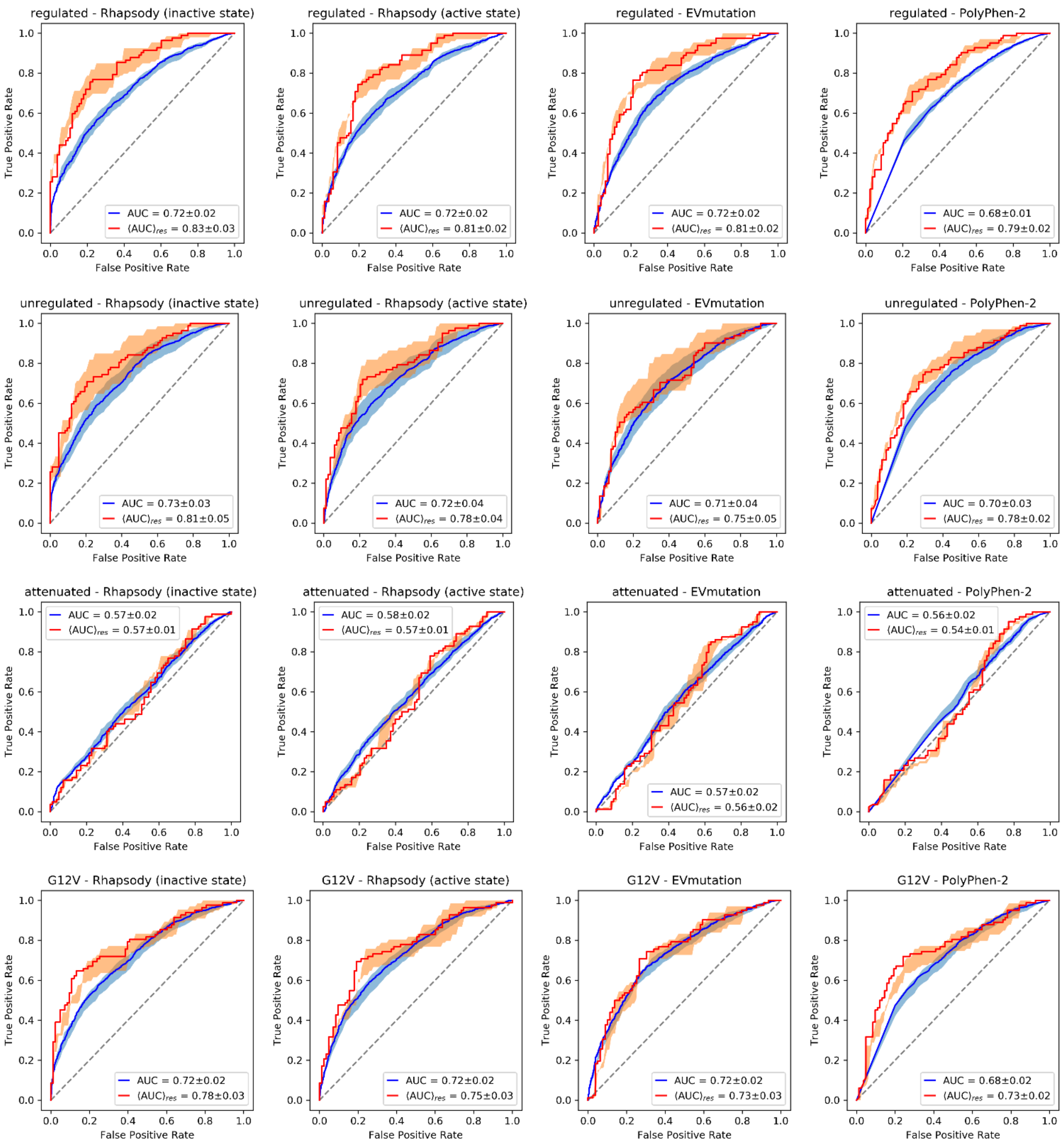


**Figure 4—figure supplement 3: ROC curves of Ras predictions**

Extended version of **Fig. 4B** with all four experimental setups reported in (Bandaru et al., 2017).

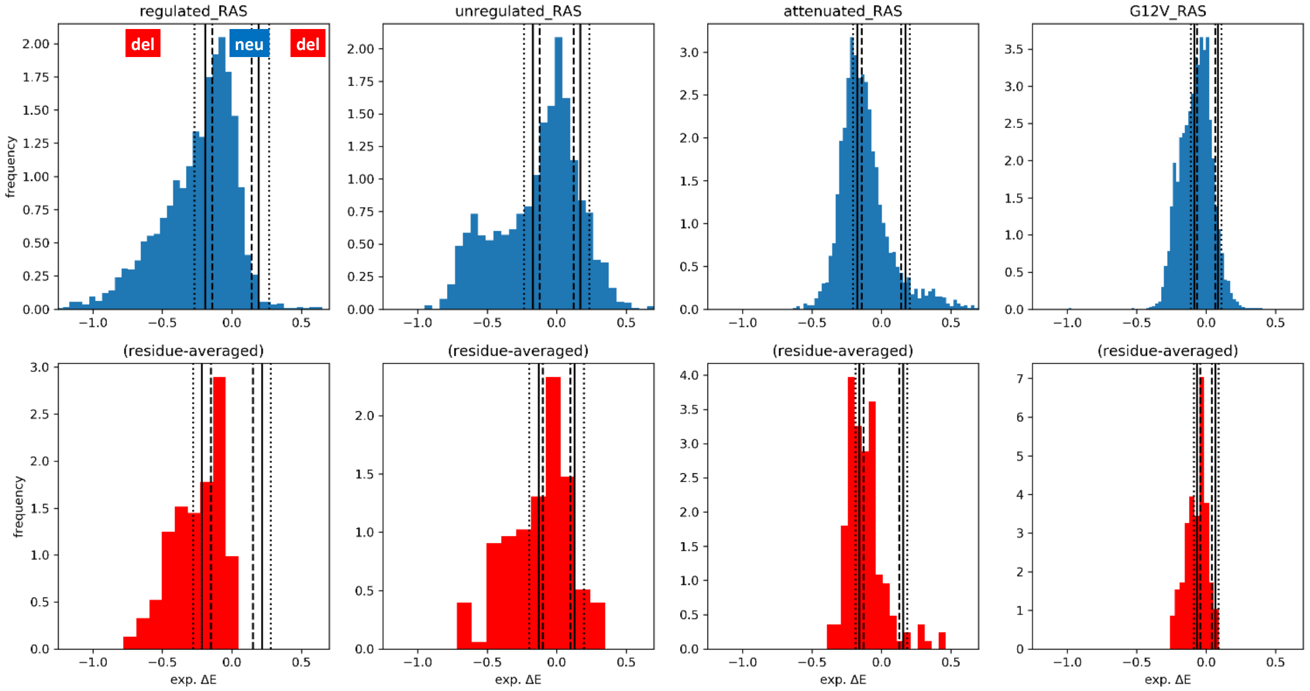


**Figure 4—figure supplement 4: Distributions of H-Ras experimental fitness scores and alternative labelling scheme**

In contrast with what presented in **Figure 4—figure supplement 2**, the dashed, solid and dotted lines mark the 40th, 50th and 60th percentiles of the *absolute* ΔE values, so that both loss- and gain-of-function variants are classified as pathogenic.

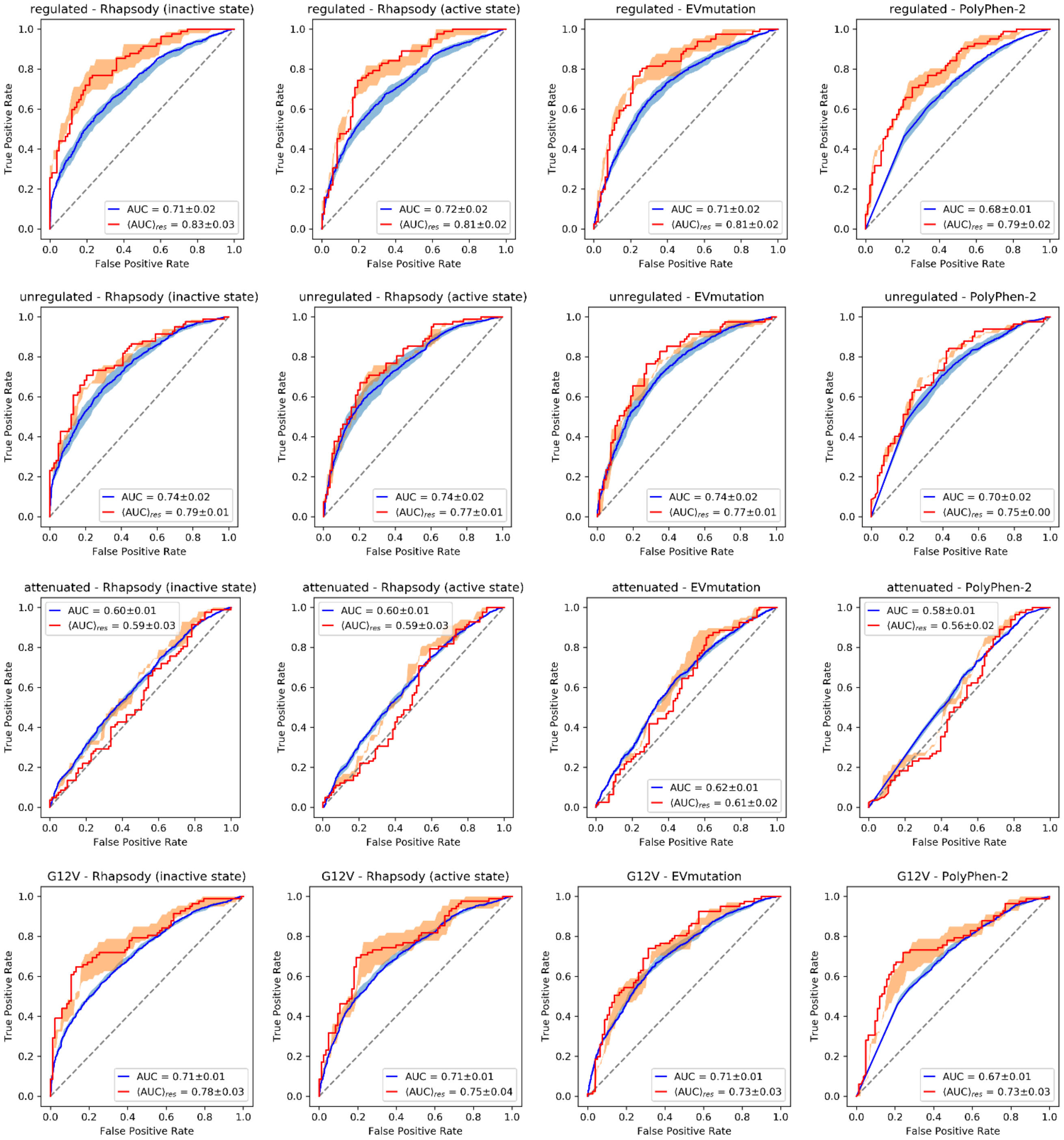


**Figure 4—figure supplement 5: ROC curves of H-Ras predictions based on alternative labelling scheme from Figure 4—figure supplement 4**

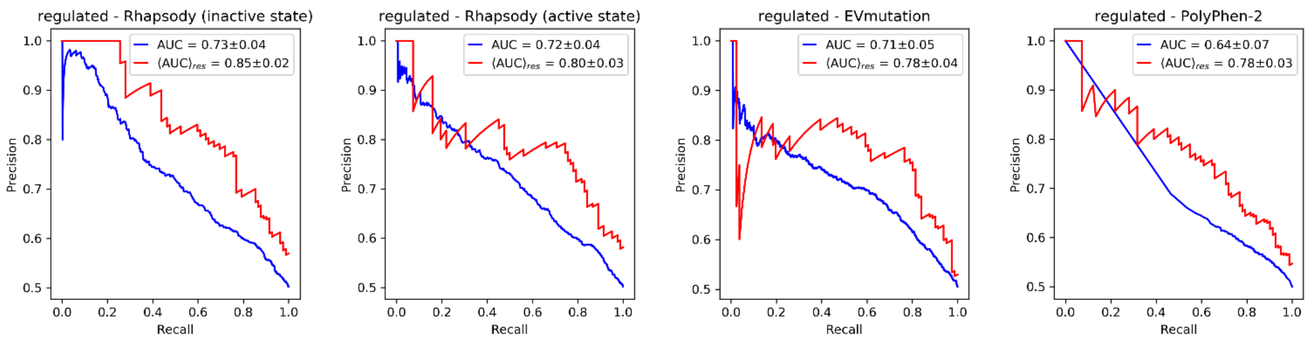


**Figure 5—figure supplement 1: Precision-Recall curves for H-Ras pathogenicity predictions**

Extended version of **Fig. 5A** with predictions from PolyPhen-2 and EVmutation.
